# Integration of the cyanophage-encoded phosphate binding protein into the cyanobacterial phosphate uptake system

**DOI:** 10.1101/2021.07.20.453049

**Authors:** Fangxin Zhao, Xingqin Lin, Kun Cai, YongLiang Jiang, Tianchi Ni, Yue Chen, Jianrong Feng, Shangyu Dang, Cong-Zhao Zhou, Qinglu Zeng

## Abstract

To acquire phosphorus, cyanobacteria use the typical bacterial ABC-type phosphate transporter, which is composed of a periplasmic high-affinity phosphate-binding protein PstS and a channel formed by two transmembrane proteins PstC and PstA. The *pstS* gene has been identified in the genomes of cyanophages that infect the unicellular cyanobacteria *Prochlorococcus* and *Synechococcus*. However, it is unknown how the cyanophage PstS interplays with the host PstC and PstA to function as a chimeric ABC transporter. Here we showed that the cyanophage P-SSM2 PstS protein was abundant in the infected *Prochlorococcus* NATL2A cells and the host phosphate uptake rate was enhanced after infection. This is consistent with our biochemical and structural analyses showing that the phage PstS protein is indeed a high-affinity phosphate-binding protein. We further modeled the complex structure of phage PstS with host PstCA and revealed three putative interfaces that may facilitate the formation of the chimeric ABC transporter. Our results provide insights into the molecular mechanism by which cyanophages enhance the phosphate uptake rate of cyanobacteria. Phosphate acquisition by infected bacteria can increase the phosphorus contents of released cellular debris and virus particles, which together constitute a significant proportion of the marine dissolved organic phosphorus pool.

## Introduction

The unicellular picocyanobacterium *Prochlorococcus* is the dominant phytoplankton in tropical and subtropical oligotrophic oceans (Partensky et al., 1999; Scanlan et al., 2009), where phosphate concentrations are in the nanomolar range and can limit primary production (Wu et al., 2000; Thingstad et al., 2005). *Prochlorococcus* has evolved several mechanisms to reduce its cellular phosphorus (P) content, including substituting non-phosphorus lipids for phospholipids (Van Mooy et al., 2006; Van Mooy et al., 2009). *Prochlorococcus* maintains an extracellular buffer of labile phosphate as a phosphate reserve to support its growth (Zubkov et al., 2015). A proton motive force is required for the import of phosphate across the outer membrane into the periplasm (Kamennaya et al., 2020). To import phosphate from the periplasm into the cytoplasm, *Prochlorococcus* cells use the high-affinity phosphate-specific transport system (Pst) and do not encode low-affinity phosphate transporters (Moore et al., 2005; Scanlan et al., 2009). The Pst system has been extensively studied in *Escherichia coli* (Lamarche et al., 2008). This ABC-type phosphate transporter comprises the periplasmic high-affinity phosphate-binding protein PstS, a channel formed by two transmembrane proteins PstC and PstA, and the intracellular homodimeric ATPase PstB (Lamarche et al., 2008; Hsieh and Wanner, 2010). In response to P limitation, the PhoR/PhoB two-component signal transduction system upregulates the transcription of phosphate-acquisition genes (Nagaya et al., 1994; Suzuki et al., 2004; Tetu et al., 2009). Consistent with the upregulation of phosphate-acquisition genes (Martiny et al., 2006; Reistetter et al., 2013), the *Prochlorococcus* strain MED4 was shown to increase its maximum uptake velocity of phosphate under P-limited conditions (Krumhardt et al., 2013). *Prochlorococcus* field populations also show enhanced phosphate uptake velocities in oceanic regions with low phosphate concentrations (Lomas et al., 2014).

Phosphate-acquisition genes have been found in the genomes of viruses (cyanophages) that infect *Prochlorococcus* and its closely related sister group marine *Synechococcus* (Sullivan et al., 2005; Sullivan et al., 2010). As lytic double-stranded DNA viruses, cyanophages comprise three morphotypes (Sullivan et al., 2003; Sabehi et al., 2012): T4-like and TIM5-like cyanomyoviruses, T7-like cyanopodoviruses, and cyanosiphoviruses. Among the 77 publically available cyanomyovirus genomes in the NCBI database (as of August 2019), 24 carry *pstS* and three carry *phoA*, an putative alkaline phosphatase gene. Due to a significant phosphorus demand for a higher nucleic acid to protein ratio, cyanophages were found to possess phosphate-acquisition genes (Jover et al., 2014). Indeed, metagenomic analysis revealed that the frequencies of P-acquisition genes in the genomes of wild cyanomyoviruses are negatively correlated with the phosphate concentrations of the marine habitats (Kelly et al., 2013), which was also found in *Prochlorococcus* genomes (Martiny et al., 2006; Martiny et al., 2009; Coleman and Chisholm, 2010).

Our previous study found that *pstS* and *phoA* genes of cyanophage S-SM1were upregulated during infection under P-limited conditions, and their expression was controlled by the host’s PhoR/PhoB system (Zeng and Chisholm, 2012). Using cyanophage P-SSM2 that contains *pstS* but not *phoA*, the transcriptomic analysis further showed that *pstS* and the adjacent gene *g247* (with unknown function) were the only two phage genes that were upregulated when *Prochlorococcus* NATL2A was infected under P-limited conditions (Lin et al., 2016). Furthermore, we discovered that under P-limited conditions the host *pstS* transcripts per cell decreased after infection, whereas the phage *pstS* transcripts were much more abundant than the host copies, resulting in more *pstS* transcripts (host and phage combined) in the infected cells (Lin et al., 2016). However, it remains unknown whether phage PstS proteins are synthesized during infection to increase the phosphate uptake rate of infected *Prochlorococcus* cells.

Here, we investigate how the cyanophage PstS protein is integrated into the phosphate acquisition system of the cyanobacterial host. Using cyanophage P-SSM2 and *Prochlorococcus* NATL2A that we have studied previously (Zeng and Chisholm, 2012; Lin et al., 2016), we compared the phosphate-binding affinities of both phage and host PstS proteins, and then measured the PstS protein abundances during infection. We also compared phosphate uptake rates of infected and uninfected *Prochlorococcus* cells. To elucidate the molecular mechanism by which the phage PstS protein binds phosphate, we solved the crystal structures of cyanophage PstS proteins. Furthermore, modeling of cyanophage PstS binding to cyanobacterial PstCA predicted several essential residues for forming a chimeric ABC transporter.

## Results

### Phosphate-binding affinities of *Prochlorococcus* and cyanophage PstS proteins

To compare the phosphate-binding affinities of PstS proteins encoded by *Prochlorococcus* NATL2A and cyanophage P-SSM2, we cloned their genes in *E. coli* and purified the recombinant PstS proteins (Supplementary Figure 1A). The binding coefficient of PstS to phosphate (the ratio of phosphate-bound PstS to the total PstS protein) showed a typical hyperbolic relationship with phosphate concentration (Figure 1A). The maximum binding coefficients (*B_max_*) for host and phage PstS proteins were 0.71 ± 0.38 and 1.12 ± 0.14, respectively. This suggested that one PstS protein binds to one phosphate molecule, which is consistent with the stoichiometry of the *E. coli* PstS protein (*B_max_* = 0.90) (Medveczky and Rosenberg, 1970; Luecke and Quiocho, 1990). The *B_max_* values of PstS proteins could be affected by the ratios of different structural conformations, with the open conformation suitable for accepting phosphate while the close conformation inaccessible to phosphate (Brautigam et al., 2014). The dissociation constants (*K*_d_) of the host and phage PstS proteins were 0.82 ± 0.44 and 1.39 ± 0.22 µM, respectively, which are comparable to those of *E*. *coli* (0.8 µM) and *Pseudomonas aeruginosa* (0.34 µM) (Medveczky and Rosenberg, 1970; Poole and Hancock, 1984). Interestingly, the *K*_d_ value of phage PstS was significantly higher than that of the host PstS (Figure 1B), indicating that the phage PstS has a relatively lower phosphate-binding affinity than the host PstS. Similarly, both cyanophages and cyanobacteria encode transaldolase genes, and the cyanophage enzymes are less efficient than the host enzymes (Thompson et al., 2011).

**Figure 1.**
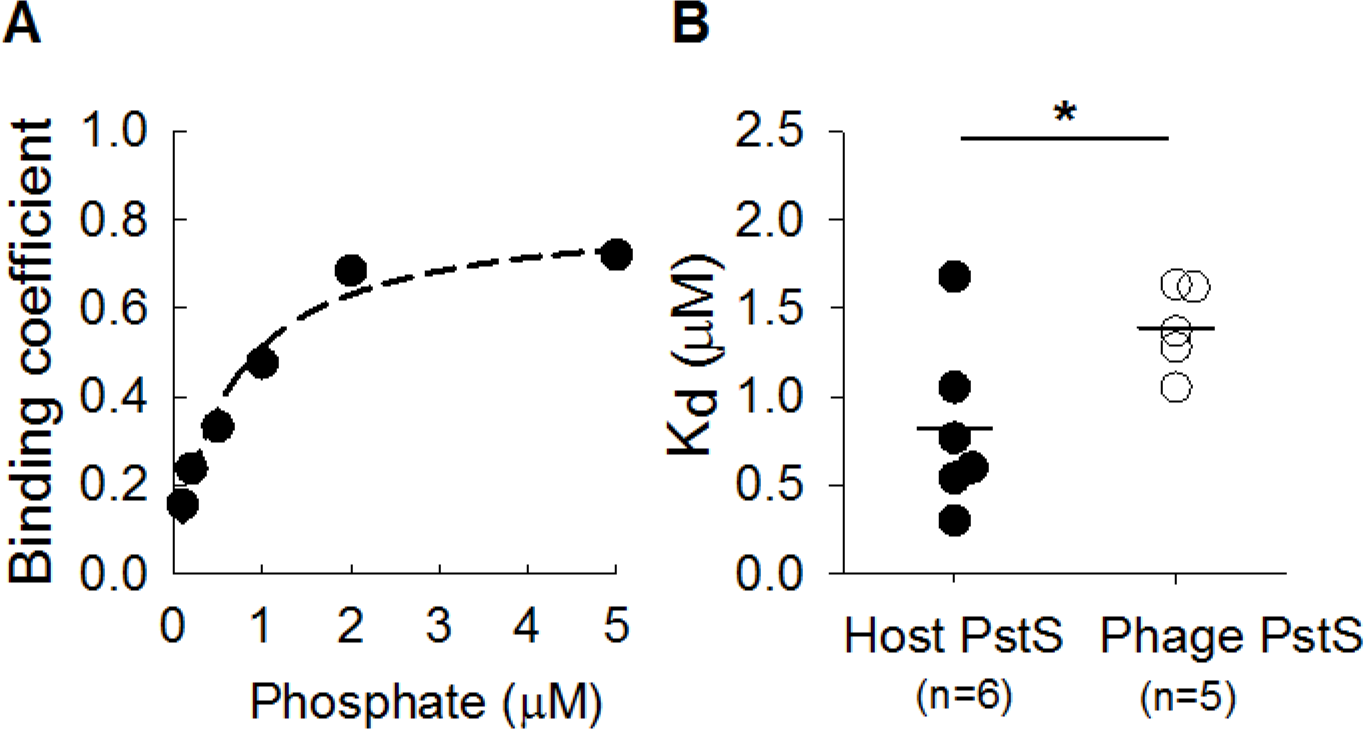
Binding coefficient and dissociation constant of PstS proteins to phosphate. **A**. A representative graph showing the binding coefficient of *Prochlorococcus* NATL2A PstS protein as a function of cold phosphate concentration. The binding coefficient is defined as the ratio of phosphate-bound PstS to the total PstS protein. A dashed line represents the non-linear regression curve fit to the Michaelis-Menten equation. **B**. The dissociation constant *K_d_* of the PstS protein to phosphate. Solid lines show the average PstS *K_d_* values of *Prochlorococcus* NATL2A (host, n = 6) and cyanophage P-SSM2 (phage, n = 5), respectively. Asterisk indicates significant difference of *K_d_* values of host and phage PstS proteins (*P* = 0.038, Student’s *t*-test).

### Host and phage PstS protein abundances during infection under P-limited conditions

We used specific antibodies (Supplementary Figure 1) to detect the host PstS protein in the uninfected cells. Similar to our previous studies (Zeng and Chisholm, 2012; Lin et al., 2016), from 24 h after resuspension in P-limited growth medium, *Prochlorococcus* NATL2A cells grew slower than those in the nutrient-replete medium (Figure 2A). Both gel electrophoresis and western blot analyses showed that the PstS protein abundance gradually increased during the progression of P limitation, while it was undetectable under nutrient-replete conditions (Supplementary Figure 2). The results are consistent with the changes of PstS protein abundances in response to P limitation in *Prochlorococcus* MED4 (Fuszard et al., 2010), *Synechococcus* WH7803 (Scanlan et al., 1993), and *Synechococcus* WH8102 (Ostrowski et al., 2010; Cox and Saito, 2013).

**Figure 2.**
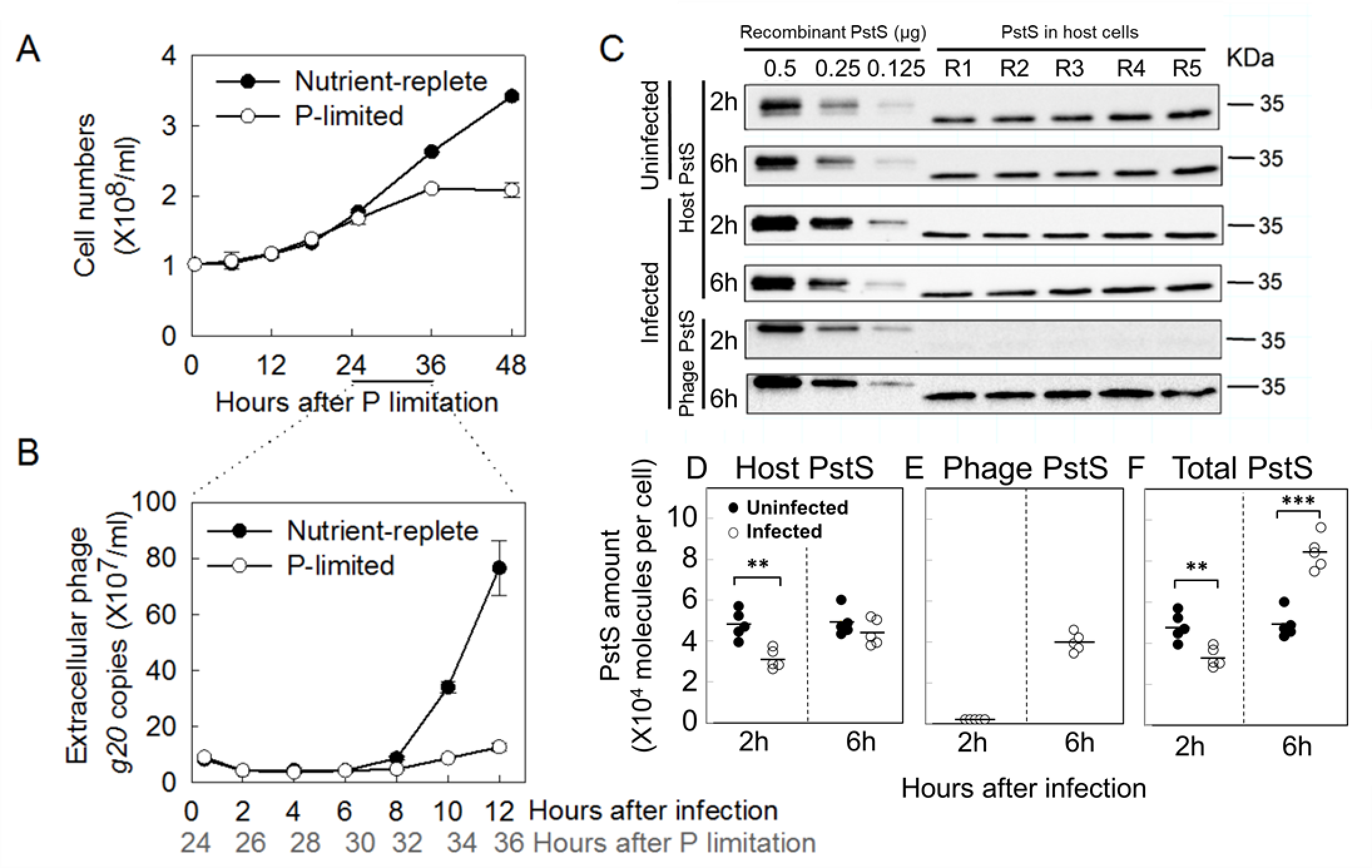
Host and phage PstS protein abundances after *Prochlorococcus* NATL2A was infected by cyanophage P-SSM2. **A**. *Prochlorococcus* NATL2A cells were spun down and resuspended in nutrient-replete or P-limited growth media**. B**. At 24 h after P limitation, *Prochlorococcus* NATL2A was infected by cyanophage P-SSM2 at a phage/host ratio of 3. Extracellular phages were measured by quantitative PCR using primers for the phage *g20* gene. Error bars in **A** and **B** indicate standard deviations from three biological replicates. **C**. Quantitative western blots of host and phage PstS proteins. At 2 h and 6 h after infection under P-limited conditions, cells were collected by centrifugation, with five biological replicates for both uninfected and infected cultures (R1 to R5). Total proteins were separated on a 12% SDS-PAGE gel (10^8^ cells per lane), transferred to a PVDF membrane, and probed with antibodies against host (top four panels) or phage PstS (bottom two panels). On the left of each gel, purified recombinant host (top four panels) or phage (bottom two panels) PstS proteins with known amounts (0.5, 0.25, 0.125 μg) were loaded as standards for protein quantification. Protein sizes are shown on the right. **D**–**F**. At 2 h and 6 h after infection under P-limited conditions, host (**D**), phage (**E**), and total (**F**) PstS proteins were quantified (Supplementary Figure 3). Error bars indicate standard deviation of five biological replicates. Asterisks indicate significant changes in the infected cells compared to the uninfected cells (** *P* < 0.005 and *** *P* < 0.0001, Student’s *t*-test).

To measure the abundances of host and phage PstS proteins, we infected *Prochlorococcus* NATL2A with cyanophage P-SSM2 (phage/host ratio = 3) at 24 h of P limitation when the host PstS protein had been highly induced (Supplementary Figure 2). Consistent with previous studies (Zeng and Chisholm, 2012; Lin et al., 2016), progeny phages were released after 8 h of infection and fewer phages were produced under P-limited conditions than those under nutrient-replete conditions (Figure 2B). Using quantitative western blotting (Figure 2C, Supplementary Figure 3), we found that under P-limited conditions the uninfected cultures had an average of 48,480 ± 6,393 PstS protein molecules per cell, which is comparable to that of *E. coli* (Medveczky and Rosenberg, 1970). During infection under P-limited conditions, the host PstS protein abundance decreased significantly by 35% at 2 h and decreased by 10% at 6 h (Figure 2D). The phage PstS protein was barely detected at 2 h after infection (Figure 2C), but increased at 6 h to 81% of the host PstS abundance in the uninfected cultures (Figure 2E). Our previous work showed that the host *pstS* transcripts decreased by 74% at 6 h after infection under P-limited conditions and the phage *pstS* transcripts were ∼5-fold higher than the host *pstS* transcripts in the uninfected cultures (Lin et al., 2016). Thus, the trends of PstS protein abundances were consistent with the transcript abundances, but with smaller changes. The variations of the transcript abundance and protein abundance of the same gene might result from translational efficiency and/or different turnover times of protein and RNA (de Sousa Abreu et al., 2009; Schwanhausser et al., 2011). Nevertheless, as a result of the high expression level of phage PstS proteins, the total PstS proteins were 71% more abundant than those in the uninfected cultures (Figure 2F).

### Phosphate uptake kinetics of *Prochlorococcus* after phage infection

Having shown that cyanophage-encoded PstS protein can bind phosphate and was expressed during infection, we wondered whether cyanophage infection affects the phosphate uptake kinetics of *Prochlorococcus* cells. We used an established ^32^P tracer method (see Materials and Methods) that has been used to measure the intracellular phosphate uptake of *Prochlorococcus* MED4 (Krumhardt et al., 2013). The phosphate uptake velocity forms a hyperbolic relationship with phosphate concentration (Supplementary Figure 4A), which is similar to a classical Michaelis-Menten curve for enzyme kinetics (Krumhardt et al., 2013). Without cyanophage infection, the maximum phosphate uptake velocity (*V*_max_) of *Prochlorococcus* NATL2A was 0.545 ± 0.096 amol phosphate cell^-1^ h^-1^ under nutrient-replete conditions and increased to 1.354 ± 0.331 amol phosphate cell^-1^ h^-1^ under P-limited conditions (Supplementary Figure 4B), which are comparable to those of *Prochlorococcus* MED4 (Krumhardt et al., 2013). The increase of *V*_max_ under P-limited conditions is consistent with the increase of PstS protein abundance (Supplementary Figure 2). The Michaelis-Menten constant (*K*_M_) of *Prochlorococcus* NATL2A was 0.357 ± 0.110 µM under nutrient-replete conditions and remained unchanged under P-limited conditions (Supplementary Figure 4C), which is within the *K*_M_ range of *Prochlorococcus* MED4 cells (Krumhardt et al., 2013).

At 6 h after infection by cyanophage P-SSM2 under P-limited conditions (Figure 3A, Supplementary Figure 4B), *V*_max_ of *Prochlorococcus* NATL2A cells increased by 57% to 2.05 amol phosphate cell^-1^ h^-1^, which is consistent with the higher amount of total PstS proteins at 6 h after infection (Figure 2F). Curiously, the phosphate uptake rate of infected cells did not decrease at 2 h (Figure 3A), although less PstS proteins were present (Figure 2D), suggesting that other proteins might function to maintain the host phosphate uptake. Because *g247* and *pstS* were the only two genes in cyanophage P-SSM2 genome that were upregulated under P-limited conditions (Lin et al., 2016), it is plausible that gp247 might play a role in phosphate acquisition during infection. A BLASTP search using gp247 protein sequence as the query did not identify any protein of known function, but two iterations of PSI-BLAST search identified several porin proteins from *Vibrio breoganii* (E value ∼10^-6^ and identity ∼30%). To test whether gp247 can form a porin structure, we predicted its 3D model using two different servers. The AI-predicted models by tFold showed a clear transporter structure formed by beta-strands (Supplementary Figure 5A). The structures predicated by the Phyre2 server also showed clear beta-strands structures, which could form a porin-like transporter in a homo-oligomeric organization (Supplementary Figure 5B). In gram-negative bacteria, porin genes have been found to be upregulated during P limitation and the expressed porins form β-barrels in the outer membrane to facilitate phosphate transport into the periplasm (Modi et al., 2015). The PstS protein can then bind phosphate and transport it across the periplasm membrane into the cytosol. Although the phage PstS protein has a higher *K*_d_ than that of the host PstS protein (Figure 1B), we did not detect significant changes of *K*_M_ after cyanophage infection, probably due to the systematic errors of this measurement (Figure 3B). Nevertheless, our results suggested that cyanophage-encoded PstS protein can enhance the host’s phosphate uptake rate during infection.

**Figure 3.**
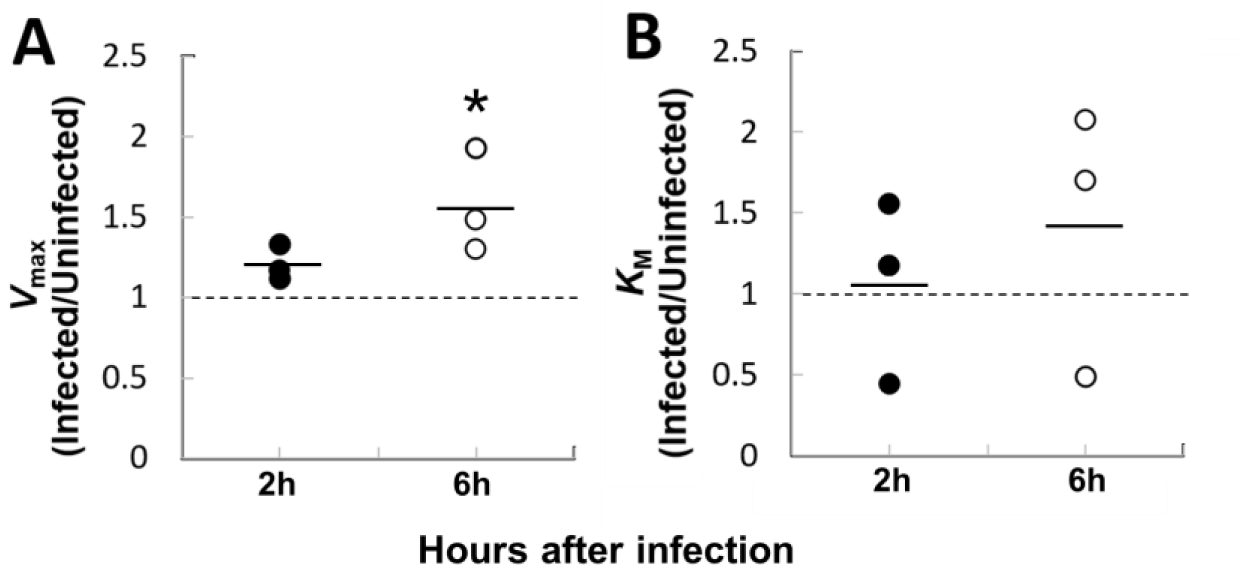
Phosphate uptake of *Prochlorococcus* NATL2A cells after infection by cyanophage P-SSM2. As in Figure 2, *Prochlorococcus* NATL2A cells were infected by cyanophage P-SSM2 at a phage/host ratio of 3 at 24 h after resuspension in P-limited growth media. At 2 h and 6 h after infection, the maximum phosphate uptake rate (*V*_max_) (**A**) and the Michaelis-Menten constant (*K*_M_) (**B**) of the infected cells were measured and normalized to those of the uninfected cells. Asterisk in **A** indicates that the normalized value is significantly larger than 1 (dashed line) (n = 3, *P* = 0.031, Student’s *t*-test).

### Structure of a cyanophage PstS protein

Currently, several structures of PstS proteins from heterotrophic bacteria have been determined (Luecke and Quiocho, 1990; Elias et al., 2012), but PstS structures of cyanobacteria and cyanophages have not been solved. To determine how PstS binds phosphate, we solved the PstS structure of cyanophage P-SSM2 at 2.25 Å resolution. To the best of our knowledge, the cyanophage PstS structure represented the first viral structure of a substrate-binding protein of an ABC-transporter. The P-SSM2 PstS structure harbors a typical “Venus flytrap” fold that is composed of two globular α/β domains held together by two β-strands (Supplementary Figure 6A). Each α/β domain contains a central mixed five-stranded β-sheet and four or five α-helices packing against the center (Supplementary Figure 6A). Similar to *E. coli* PstS (Luecke and Quiocho, 1990), one phosphate molecule binds to the cleft between the two α/β domains (Supplementary Figure 6A). Moreover, the phosphate-binding residues Ser30, Ser59, Asp77, Arg146, Asp148, and Ser150 (Supplementary Figure 6B) are highly conserved among the PstS proteins with solved structures (Supplementary Figure 7).

### Modeling the interactions between cyanophage PstS and host PstCA

To be functional in the infected host cells, the cyanophage PstS protein needs to interact with the host PstC and PstA proteins to form a chimeric ABC transporter for phosphate uptake, since cyanophage genomes do not contain *pstC* and *pstA* genes. To investigate the interaction surface of PstS with the transmembrane PstCA complex, we tried to express and purify *Prochlorococcus* PstC and PstA proteins in *E. coli*, but could not obtain soluble proteins. Hence, we simulated the PstCA structure of *Prochlorococcus* NATL2A and docked P-SSM2 PstS onto the host PstCA complex (see Materials and Methods). Our simulated model showed that the phage PstS positions above the extracellular face of PstCA with its phosphate-binding pocket facing towards PstCA (Figure 4A). PstS interacts with the host PstCA complex mainly via three interfaces, an α-helix α2 (interface 1), a η-helix η1 and flanking loops (interface 2), and a loop L_α4-β7_ (interface 3) (Figure 4A). Detailed analysis showed that Ser61, Lys64 and Asp68 of interface 1, Lys87 of interface 2, and Thr171, Lys173, and Ala176 of interface 3 contribute to the major polar interactions with PstCA (Figure 4B). Specifically, the interface 1 interacts with Phe163, Asn255, Asn256 and Glu278 of PstC, and the interface 2 interacts with Glu143 and Arg147 of PstC and Glu263 of PstA (Figure 4B). In addition, the interface 3 interacts with Tyr258, Asn259, and Tyr265 of PstA (Figure 4B). Therefore, our results suggested that the phage PstS protein can interact with the host PstCA complex to form a chimeric ABC transporter, which provides a molecular mechanism by which cyanophage infection enhances the phosphate uptake rate of cyanobacteria (Figure 3A).

**Figure 4.**
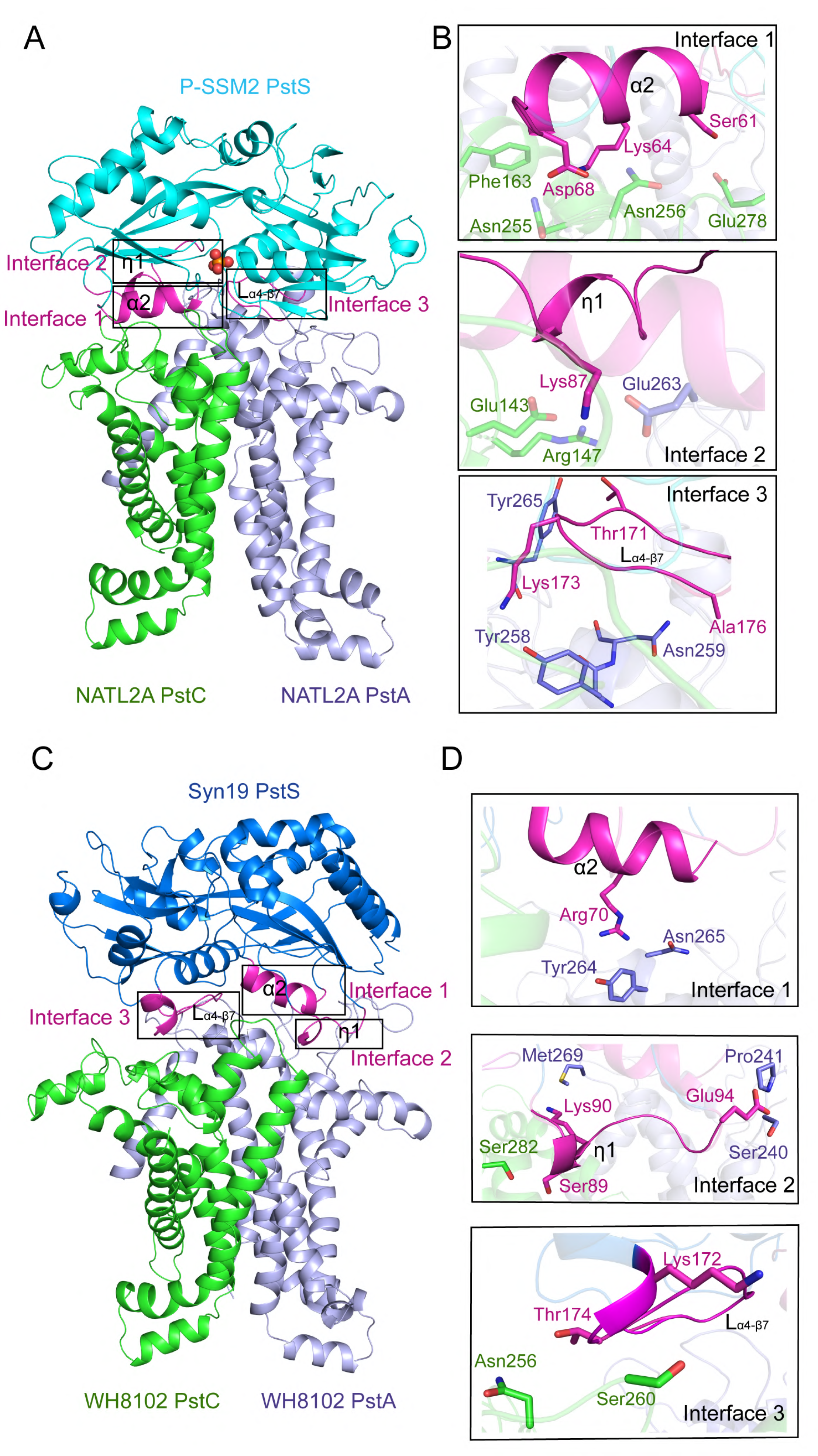
Simulated models of cyanophage PstS binding to the host PstCA complex. **A**. A simulated model shows the interaction of the PstS protein of cyanophage P-SSM2 (cyan) with PstC (green) and PstA (lightblue) of *Prochlorococcus* NATL2A. The three interface regions of PstS that interact with the PstCA complex are shown in purple and highlighted by boxes. **B**. The detailed interaction networks of the three interface regions of P-SSM2 PstS. Residues involved in the interactions are shown in sticks and are colored in purple for PstS, green for PstC and lightblue for PstA. **C.** A simulated model of PstS protein of cyanophage Syn19 (blue) interacting with PstC (green) and PstA (lightblue) of *Synechococcus* WH8102. Comparing to P-SSM2 PstS, the three interface regions of Syn19 PstS (purple) showed a ∼180° rotation against the host PstCA complex. **D.** The detailed interacting networks of the three interface regions of Syn19 PstS binding to PstCA. The color scheme in **D** is the same as in **B**.

### Two groups of cyanophage PstS proteins with different interface sequences

To investigate whether the three interface sequences are conserved in cyanophage PstS proteins, we built a maximum-likelihood phylogenetic tree using the PstS protein sequences from the currently available cyanophage genomes, together with their host PstS sequences. Based on the tree, we grouped the PstS sequences into four groups, termed I, II, III, and SphX (Figure 5). The group I PstS contained *Prochlorococcus*, *Synechococcus*, and cyanophage sequences, each forming a separate clade (Figure 5A). PstS of cyanophage P-SSM2 is in the cyanophage clade of group I PstS sequences. The group II PstS contained cyanophage sequences and the group III PstS contained *Synechococcus* sequences (Figure 5A). The SphX group contained *Synechococcus* sequences that are closely related to the phosphate-binding protein SphX of the freshwater cyanobacterium *Synechococcus* PCC7942 (Scanlan et al., 2009) (Figure 5). Group I cyanophage PstS sequences fell into a phylogenetic clade within cyanobacterial sequences, while group II cyanophage sequences formed a distinct clade (Figure 5A), suggesting that cyanophages might have gained the *pstS* gene from their cyanobacterial hosts in at least two separate evolutionary events. Sequence alignment showed that the three interface regions of P-SSM2 PstS are highly conserved among group I PstS sequences, but are quite different from the corresponding regions in the groups II, III and SphX sequences (Figure 5B). Since the host PstC and PstA residues that interact with the group I PstS of cyanophage P-SSM2 are highly conserved in the sequenced *Prochlorococcus* and *Synechococcus* genomes (Supplementary Figure 8) (see Discussion), the group II cyanophage PstS proteins may interact with the host PstCA complex in a different way compared to the group I PstS proteins.

**Figure 5.**
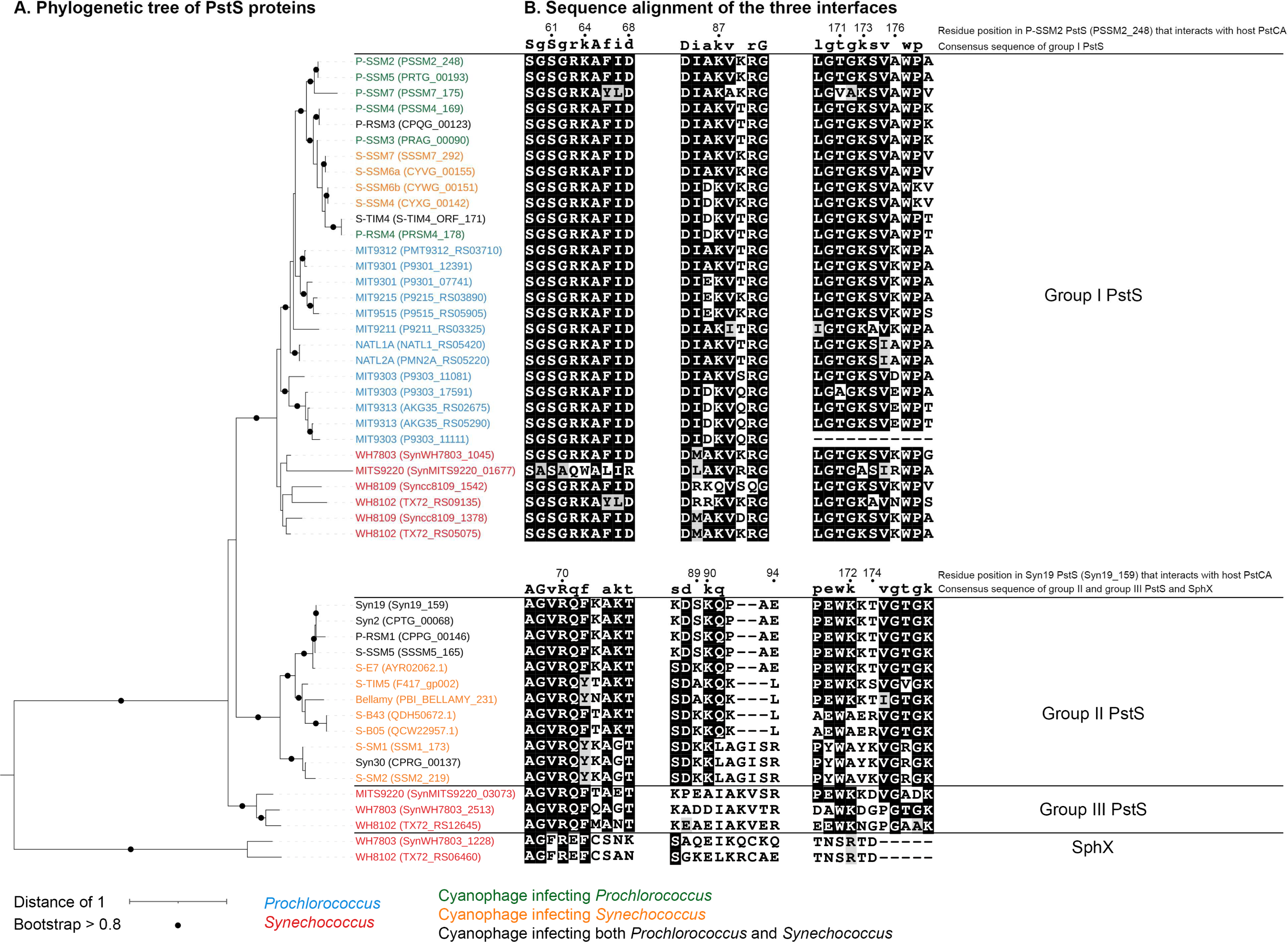
Phylogeny of the PstS sequences from cyanobacteria and cyanophages. **A**. Phylogenetic tree of the PstS protein sequences built using the maximum likelihood method. The PstS sequence of *Pseudomonas aeruginosa* was used as an outgroup to re-root the tree (not shown). Locus_tag or accession number of each protein is shown in parentheses. In the tree, PstS protein sequences form four groups: I, II, III, and SphX. **B**. Sequence alignments of the three interface regions of group I PstS proteins (top), and groups II, III, and SphX proteins (bottom). The numbers above the top and bottom alignments indicate the amino acid residue positions of P-SSM2 and Syn19 PstS proteins, respectively. Degree of conservation is indicated with background shading, with dark for strongly conserved and light for moderately conserved residues.

To explore how the group II cyanophage PstS protein interacts with the host PstCA complex, we solved the crystal structure of a group II PstS protein from cyanophage Syn19 at 1.70 Å resolution (Supplementary Figure 9B). The P-SSM2 and Syn19 PstS proteins share a high structural similarity to each other with a root-mean square deviation (RMSD) value of 0.80 Å over 255 Cα atoms (Supplementary Figure 9D). We then simulated the PstCA structure of *Synechococcus* WH8102, which is the original host of Syn19, and docked Syn19 PstS onto the host PstCA complex (Figure 4C). Compared with the interaction model of P-SSM2 PstS and *Prochlorococcus* NATL2A PstCA (Figure 4A), Syn19 PstS also harbors three interfaces that are structurally similar to those of P-SSM2 PstS (Supplementary Figure 9D). However, when the PstCA structures of *Prochlorococcus* NATL2A and *Synechococcus* WH8102 were superimposed, the Syn19 PstS shows a ∼180° rotation as compared to P-SSM2 PstS (compare Figures 4A and 4C). As a result, the interfaces 1 and 2 of Syn19 interact with WH8102 PstA whereas the interface 3 binds to WH8102 PstC (Figure 4C). On the contrary, the interfaces 1 and 2 of P-SSM2 PstS interact with NATL2A PstC, and the interface 3 binds to NATL2A PstA (Figure 4A). The residues Arg70 of interface 1, Ser89, Lys90, Glu94 of interface 2, and Lys172 and Thr174 of interface 3 contribute to the interaction with WH8102 PstCA (Figure 4D). Sequence alignment showed that most of the interface residues are conserved among group II cyanophage PstS proteins (Figure 5B). Taken together, our results suggested that both group I and II cyanophage PstS proteins could recognize the host PstCA complex to organize into a functional phosphate transporter that enhances the phosphate uptake rate of infected host cells.

## Discussion

In this study, we showed that the cyanophage P-SSM2 PstS protein is a functional phosphate-binding protein and is abundantly expressed during infection under P-limited conditions, resulting in more PstS proteins in the infected cultures than in the uninfected cultures. Consistently, the maximum phosphate uptake velocity of infected *Prochlorococcus* NATL2A cultures was higher than that of the uninfected cultures. The high-resolution crystal structures of cyanophage PstS proteins revealed key phosphate-binding residues that are conserved in bacterial and cyanophage PstS proteins. By docking the cyanophage PstS structure onto the simulated structure of the host PstCA complex, we were able to predict essential residues for the interaction of cyanophage PstS with the host PstCA complex, suggesting the formation of chimeric ABC transporters in the infected host cells. By using a combination of enzymatic, biochemical, and structural analyses, our work provides molecular mechanisms by which cyanophage PstS protein is integrated into the phosphate uptake system of the cyanobacterial host cells.

The phosphate ABC transporter is composed of PstS, the transmembrane channel proteins PstC and PstA, and the ATPase PstB (Lamarche et al., 2008; Hsieh and Wanner, 2010), but the currently available cyanophage genomes only contain the *pstS* gene, lacking the other three components. For ABC transporters to achieve maximal import activity, mathematic models indicate that the concentrations of substrate-binding proteins (e.g., PstS) should be higher than those of the transporters and substrates (Bosdriesz et al., 2015). A higher concentration of the substrate-binding protein increases the encounter rate of the transporter with the substrate and thus increases the substrate uptake rate of the transporter (Ames and Lever, 1970; Bosdriesz et al., 2015). Therefore, PstS protein abundance is the rate-limiting step for phosphate uptake and this may explain why cyanophages express additional PstS proteins during infection (Figure 2E). Enhancing the phosphate uptake velocity of infected host cells (Figure 3A) can fulfil the high phosphorus demand of cyanophages (Jover et al., 2014), which may confer cyanophages a selective advantage under P-limited oceanic regions (Kelly et al., 2013).

Our phylogenetic analysis identified four groups of PstS proteins in cyanobacterial/cyanophage genomes. *Prochlorococcus* and marine *Synechococcus* genomes all contain at least one copy of the group I *pstS* gene, and some contain additional *pstS*/*sphX* genes (Figure 5A). For example, the genome of *Synechococcus* sp. WH8102 encodes two group I *pstS* genes, one group III *pstS* gene, and one *sphX* gene (Figure 5A). It was hypothesized that different *pstS*/*sphX* genes may encode proteins with different phosphate-binding affinities that could be used in different environmental conditions (Scanlan et al., 2009). Despite multiple *pstS*/*sphX* genes, each cyanobacterial genome only encodes one *pstA* gene and one *pstC* gene (Martiny et al., 2006). Therefore, different PstS/SphX proteins in a cyanobacterial cell should be able to interact with the same PstCA complex. Indeed, our structural simulations suggested that both groups I and II cyanophage PstS proteins are able to interact with the host PstCA complex, although using different interface residues (Figure 4) that are conserved within each group (Figure 5B). Future site-directed mutagenesis and structural analysis are needed to verify the interface residues we predicted in this study.

Nutrient acquisition by infected cells affects the elemental stoichiometry of released materials after cell lysis (Jover et al., 2014). The dissolved organic phosphorus (DOP) released after infection comprises cellular debris and virus particles, the total amount of which depends on both the phosphorus content of uninfected host cells and the newly acquired phosphorus during infection. We showed that the maximum phosphate uptake velocity of infected *Prochlorococcus* cells increased by 57% at 6 h after infection (Figure 3A), indicating that more phosphate is acquired by the infected cells and thus more DOP should be released after infection. Marine virus particles have been estimated to constitute >5% of the total DOP pool in the surface waters of several oceanic regions (Jover et al., 2014) and cellular debris should constitute an even large proportion in the marine DOP pool. Thus, by manipulating the host phosphate ABC transporter system, cyanophages have the potential to affect phosphorus cycling in the oceans.

## Acknowledgements

This study is supported by grants to Qinglu Zeng from the Hong Kong Branch of Southern Marine Science and Engineering Guangdong Laboratory (Guangzhou) (Project numbers SMSEGL20SC01 and GML2019ZD0409). We thank Haiwei Luo for helpful discussion.

## Author Contributions

Conceptualization, Q.Z.; Methodology, Q.Z., F.Z., X.L., K.C., Y.J., T.N., and C-Z.Z.; Formal analysis, X.L., Y.J, Y.C., and F.Z.; Investigation, Q.Z., F.Z., X.L., K.C., Y.J., T.N, Y.C., J.F., S.D., C-Z.Z.; Writing – Original Draft, Q.Z., X.L., Y.J., F.Z.; Writing – Review & Editing, all authors.

## Materials and methods

### Expression and purification of recombinant PstS proteins

The *pstS* genes of *Prochlorococcus* NATL2A, cyanophage P-SSM2, and cyanophage Syn19 were amplified by PCR using primers listed in Supplementary Table 1. PCR products were cloned into the pET-22b vector, and then transformed into *Escherichia coli* BL21 (DE3) cells harboring the pKY206 plasmid. *E. coli* cells were grown in 1 L LB medium (10 g Bacto tryptone, 10 g NaCl, and 5 g yeast extract per liter) with 50 µg ml^-1^ ampicillin and 5 µg ml^-1^ tetracycline at 37°C for 5 h until OD_600_ = 0.8. Recombinant proteins with an C-terminal hexahistidine tag were induced with 0.2 mM IPTG (isopropyl β-D-1-thiogalactopyranoside) for 20 h at 16°C. *E. coli* cells were harvested by centrifugation at 8,000 g for 10 min and resuspended in the lysis buffer (20 mM Tris-HCl, 200 mM NaCl, 5% glycerol, 5 mM sodium phosphate, pH 7.5). After sonication for 30 min, the cultures were spun down at 12,000 g for 30 min and the supernatants were loaded onto a His-Select Nickel Affinity gel (GE Healthcare). Recombinant PstS proteins were eluted with the elution buffer (500 mM imidazole, 20 mM Tris-HCl, 200 mM NaCl, 5% glycerol, 5 mM sodium phosphate, pH 7.5) and then further purified using HiLoad 16/60 Superdex 200 columns (GE Healthcare). For crystallization, PstS proteins with bound phosphate substrates were purified using HiLoad 16/60 Superdex 200 columns pre-equilibrated with 20 mM Tris-HCl, 200 mM NaCl, 5% glycerol, 5 mM sodium phosphate, 14 mM β-mercaptoethanol, pH 7.5. For phosphate binding affinity assays, PstS proteins without bound phosphate substrates were purified using columns pre-equilibrated with 20 mM Tris-HCl, 200 mM NaCl, 5% glycerol, pH 7.5.

### Measurement of phosphate-binding affinity of recombinant PstS proteins

Prior to the measurement of phosphate-binding affinity, we removed the residual phosphate substrates from the purified PstS proteins by dialyzing in the Tris buffer (20 mM Tris-HCl, 200 mM NaCl, pH 7.5) at 4°C for 24 h using the Slide-A-Lyzer mini dialysis devices (20K MWCO, Thermo Fisher Scientific). After dialysis, protein concentrations were measured using the DC Protein Assay Kit (Bio-Rad) with BSA (bovine serum albumin) as standards.

Equilibrium dialysis was performed to determine the dissociation constant (*K*_d_) of the recombinant PstS proteins (Poole and Hancock, 1984). In each Slide-A-Lyzer mini dialysis unit (20K), 4 µg protein was placed in the top dialysis chamber, and Tris buffer containing trace amount of ^32^P-labeled orthophosphoric acid (∼1.2 pmol and ∼1 µCi, PerkinElmer) and different concentrations of cold phosphate (NaH_2_PO_4_) was placed in the bottom dialysis buffer chamber. After shaking for 24 h at room temperature, 100 µl samples were taken from the dialysis chamber and the dialysis buffer chamber, respectively. Radioactivity was measured by adding the samples into 4 ml liquid scintillation cocktail (OptiphaseHiSafe 3, PerkinElmer) and counting with a liquid scintillation counter (Wallac Win Spectral 1414, PerkinElmer). The dissociation constant *K*_d_ was determined using the following equation (Michaelis et al., 2011; Viaene et al., 2013):

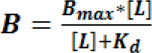

where *B* is the binding coefficient, *B*_max_ is the maximum binding coefficient, and *L* is the concentration of free phosphate. Non-linear regression fitting was performed using SigmaPlot v12.5 (Systat Software).

### Infection of *Prochlorococcus* NATL2A by cyanophage P-SSM2

Infection of *Prochlorococcus* NATL2A by cyanophage P-SSM2 under P-limited conditions was carried out as we described previously (Zeng and Chisholm, 2012; Lin et al., 2016). The axenic *Prochlorococcus* NATL2A culture was maintained at 24°C under constant cool white light (∼30 µE m^-2^ s^-1^) in the Pro99 growth medium (Moore et al., 2002) that is based on Port Shelter seawater from Hong Kong. Fresh cyanophage P-SSM2 lysate was concentrated with Amicon Ultra-15 30K Centrifugal Filter Units (Millipore) at 3,000 g for 15 min, washed twice with sterile seawater, and resuspended in the same medium. Prior to infection, mid-log *Prochlorococcus* NATL2A cultures were centrifuged at 10,000 g for 15 min at 21°C, washed with the nutrient-replete Pro99 medium (with 50 μM phosphate) or the P-depleted Pro99 medium (without added phosphate), and resuspended in the same media. After 24 h of resuspension, *Prochlorococcus* NATL2A cells were mixed with cyanophage P-SSM2 at a phage/host ratio of 3. Cell numbers were measured by flow cytometry (BD FACSCalibur, BD Biosciences). Extracellular phages were measured by quantitative PCR using primers for *g20* (Supplementary Table 1).

### Quantification of host and phage PstS proteins using specific antibodies

The purified recombinant PstS proteins of *Prochlorococcus* NATL2A and cyanophage P-SSM2 were used as antigens to generate antibodies (Antibody host: rabbit; custom ordered from MW Biotech). The specificity of antibodies was confirmed using recombinant PstS proteins (Supplementary Figure 1) and *Prochlorococcus* NATL2A cells (Figure 2C).

To detect PstS proteins by western blot, purified proteins or total proteins of *Prochlorococcus* cultures were denatured at 95°C for 15 min in the loading buffer (62.5 mM Tris-Cl, pH 6.8, 2% SDS, 0.05% bromophenol blue, 1% glycerol, and 0.05% β-mercaptoethanol). Denatured proteins were separated on a 12% SDS-PAGE gel, stained with Coomassie Blue, and visualized with the ChemiDoc Imaging System (Bio-Rad). For western blotting, proteins were transferred from the SDS-PAGE gel (without staining) onto a PVDF membrane and probed with primary antibodies against *Prochlorococcus* NATL2A PstS or cyanophage P-SSM2 PstS. The membrane was then incubated with an HRP-conjugated secondary antibody (ECL anti-rabbit IgG, GE Healthcare) and PstS bands were visualized with the ChemiDoc Imaging System (Bio-Rad).

For absolute quantification of PstS proteins in *Prochlorococcus* NATL2A cells, total proteins from 10^8^ cells were separated on 12% SDS-PAGE alongside serial dilutions of purified recombinant PstS proteins with known amounts. Proteins were then transferred to a PVDF membrane for western blot analysis using PstS antibodies (Figure 2C). A standard curve was generated using the signal volume of recombinant PstS bands and the corresponding protein amounts (Supplementary Figure 3A). Based on the standard curve, the average number of PstS protein molecules per cell was calculated (Supplementary Figure 3B).

### Phosphate uptake kinetics of *Prochlorococcus* cells

Phosphate uptake kinetics of *Prochlorococcus* NATL2A cells was measured following an established method that has been used for *Prochlorococcus* MED4 (Krumhardt et al., 2013). Briefly, 12 ml culture was centrifuged at 10,000 g for 15 min at 21°C and resuspended with the same volume of the Pro99 medium without addition of phosphate. After resuspension, aliquots of 1 ml cultures were transferred to clear Eppendorf tubes, which contained trace amount of ^32^P-labeled orthophosphoric acid (∼1 µCi, Perkin Elmer) and different concentrations of cold PO_4_ (from 0.02 μM to 20 μM). Cultures were incubated for 60 min at 24°C at a light level of ∼30 µE m^-2^ s^-1^ to allow linear uptake of phosphate (Supplementary Figure 10). Cultures were then filtered at a vacuum pressure of ∼100 mm Hg through a 0.22 µm polycarbonate filter that was supported by a Whatman GF/F filter. Prior to filtration, the filters were pre-soaked with the Pro99 medium amended with 0.5 mM PO_4_ to minimize non-specific adhesion of ^32^P on to the filters. After filtration, the filters were soaked in a basic oxalate reagent for 10 min and dried by filtration for 30 sec. Since oxalate removes the extracellular phosphate buffer of cyanobacterial cells (Zubkov et al., 2015), the remaining ^32^P reflected intracellular phosphate uptake by *Prochlorococcus* cells. The filters were immersed into 4 ml liquid scintillation cocktail (Optiphase HiSafe 3, Perkin Elmer) and the radioactivity of each filter was measured by a liquid scintillation counter (Wallac Win Spectral 1414, PerkinElmer). A filter without any cells was measured as a blank control to reflect the background ^32^P level.

The phosphate uptake velocity (*V*) of *Prochlorococcus* NATL2A was determined by the following equation (Fu et al., 2005; Krumhardt et al., 2013):

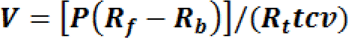

where *R*_f_ is the ^32^P radioactivity on the filters with cells, *R*_b_ denotes the radioactivity on the blank control filter without cells, and *R*_t_ is the total radioactivity in the 1 ml culture. *P* is the concentration of cold PO_4_ (amol per liter), while *t*, *c* and *v* represents incubation time (h), cell concentration (cells per liter) and volume of the filtered culture (liter), respectively. The velocity (amol cell^-1^ h^-1^) was plotted against cold phosphate concentration and the curve was fitted to the Michaelis-Menten equation:

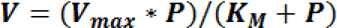

where *V*_max_ represents the maximum velocity of phosphate uptake and *K*_M_ represents the Michaelis-Menten constant. The non-linear regression was performed on SigmaPlotv12.5 (Systat Software, USA).

### Crystallization of PstS with phosphate

Fractions containing the target protein were pooled and concentrated to 8 mg ml^-1^ for crystallization. Crystals were grown by mixing the protein sample with the reservoir solution at a 1:1 ratio in hanging drops at 13°C. The crystallization buffer was composed of 20 mM Tris-HCl, pH 7.5, 200 mM NaCl, 5% glycerol, 5 mM sodium phosphate buffer, pH 7.5, 1 mM DTT. Crystals of PstS from P-SSM2 appeared in the condition containing 25% (w/v) polyethylene glycol 3350, 0.1 M citric acid, pH 3.5, while the Crystals of PstS from Syn19 appeared in the condition containing 18% (w/v) polyethylene glycol 3350, 0.2 M sodium formate.

### Data Collection and structure determination

For diffraction analysis, the crystals were pooled and flash-frozen in liquid nitrogen after soaking in the glycerol cryoprotectant. X-ray diffraction data were collected at the beamline BL17U at the Shanghai Synchrotron Radiation Facility (SSRF). Diffraction images were processed and scaled with HKL-2000 program package (Otwinowski and Minor, 1997) to the highest resolutions of 2.25 Å for P-SSM2 PstS and 1.70 Å for Syn19 PstS, respectively. The structures were determined by molecular replacement using the program Molrep of the CCP4 (Collaborative Computational Project, 1994) with *E. coli* PstS (PDB accession code 1IXH) as the search template (Wang et al., 1997). The structure refinement was performed by using the programs Coot (Emsley and Cowtan, 2004) and Refmac. The quality of the structures was analyzed by MolProbity (Chen et al., 2010). The parameters of crystal data collection and structure refinement for P-SSM2 and Syn19 PstS proteins are listed in Supplementary Tables 2 and 3, respectively. All figures showing structures were prepared with PyMOL.

### Modeling the interaction of cyanophage PstS with the host PstCA complex

We simulated the PstCA structure of *Prochlorococcus* NATL2A by SWISS-MODEL (https://swissmodel.expasy.org/) using the published structure of the maltose ABC transporter MalFG (Oldham et al., 2007). Then, we docked the phage PstS structure onto the simulated PstCA models using HADDOCK (http://haddock.science.uu.nl/services/HADDOCK2.2). HADDOCK clustered 103 structures in 11 clusters, which represent 51.5 % of the water-refined models HADDOCK generated. The best cluster, which is the most reliable according to HADDOCK, possess a Haddock score of -136.9(+/-8.7), Z-score of -2.2 and a RMSD value of 0.6 (+/-0.3) Å. The Z-score indicates how many standard deviations from the average this cluster is located in terms of score (the more negative the better).

### Phylogenetic analysis

Among the 77 cyanomyovirus genomes available in the NCBI database (as of August 2019), PstS protein sequences were identified in 24 cyanomyovirus genomes. Those 24 cyanomyoviruses were shown to infect 9 *Prochlorococcus* strains and 4 *Synechococcus* strains (Sullivan et al., 2003; Sullivan et al., 2010; Sabehi et al., 2012; Hua et al., 2017; Enav et al., 2018; Zborowsky and Lindell, 2019; Jiang et al., 2020; Wang et al., 2020). PstS protein sequences of the 24 cyanomyoviruses and 13 cyanobacterial host strains were downloaded from NCBI. For phylogenetic analysis, PstS protein sequences were aligned using Clustal Omega (Madeira et al., 2019) and visualized by BOXSHADE (https://embnet.vital-it.ch/software/BOX_form.html). Phylogenetic inference was based on the resulting alignment and conducted using the RAxML software (Stamatakis, 2014). The phylogenetic tree was visualized by Interactive Tree of Life (https://itol.embl.de/) (Letunic and Bork, 2016).

### Data Availability

PstS structures of P-SSM2 and Syn19 have been deposited in the Protein Data Bank (PDB) with the accession numbers 7DYP and 7DYO, respectively.

---

**Supplementary Figure 1.**
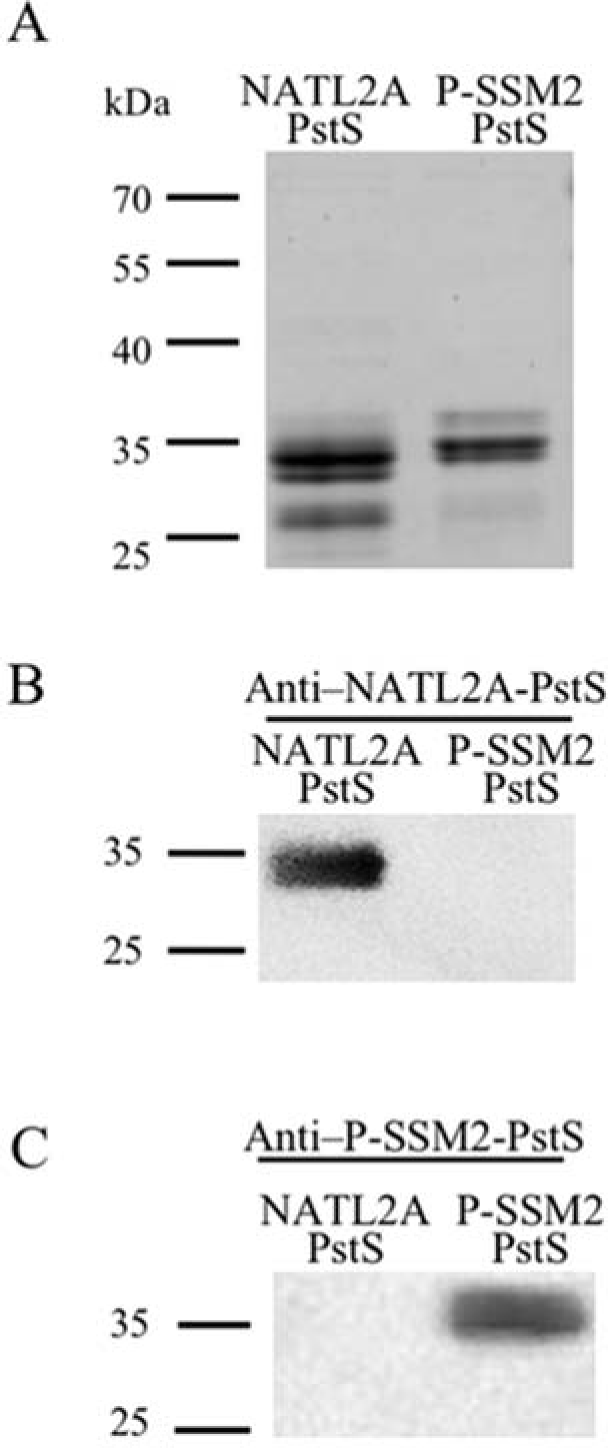
Specificity of antibodies against the PstS proteins of *Prochlorococcus* NATL2A and cyanophage P-SSM2. The His-tagged recombinant PstS proteins of *Prochlorococcus* NATL2A and cyanophage P-SSM2 were purified and separated on 12% SDS-PAGE. Proteins were stained with Coomassie Blue (**A**) or transferred to membranes and probed using antibodies against NATL2A PstS (**B**) and P-SSM2 PstS (**C**). In **B** and **C**, 0.5 µg protein was loaded in each lane. Protein sizes are shown on the left of each gel.

**Supplementary Figure 2.**
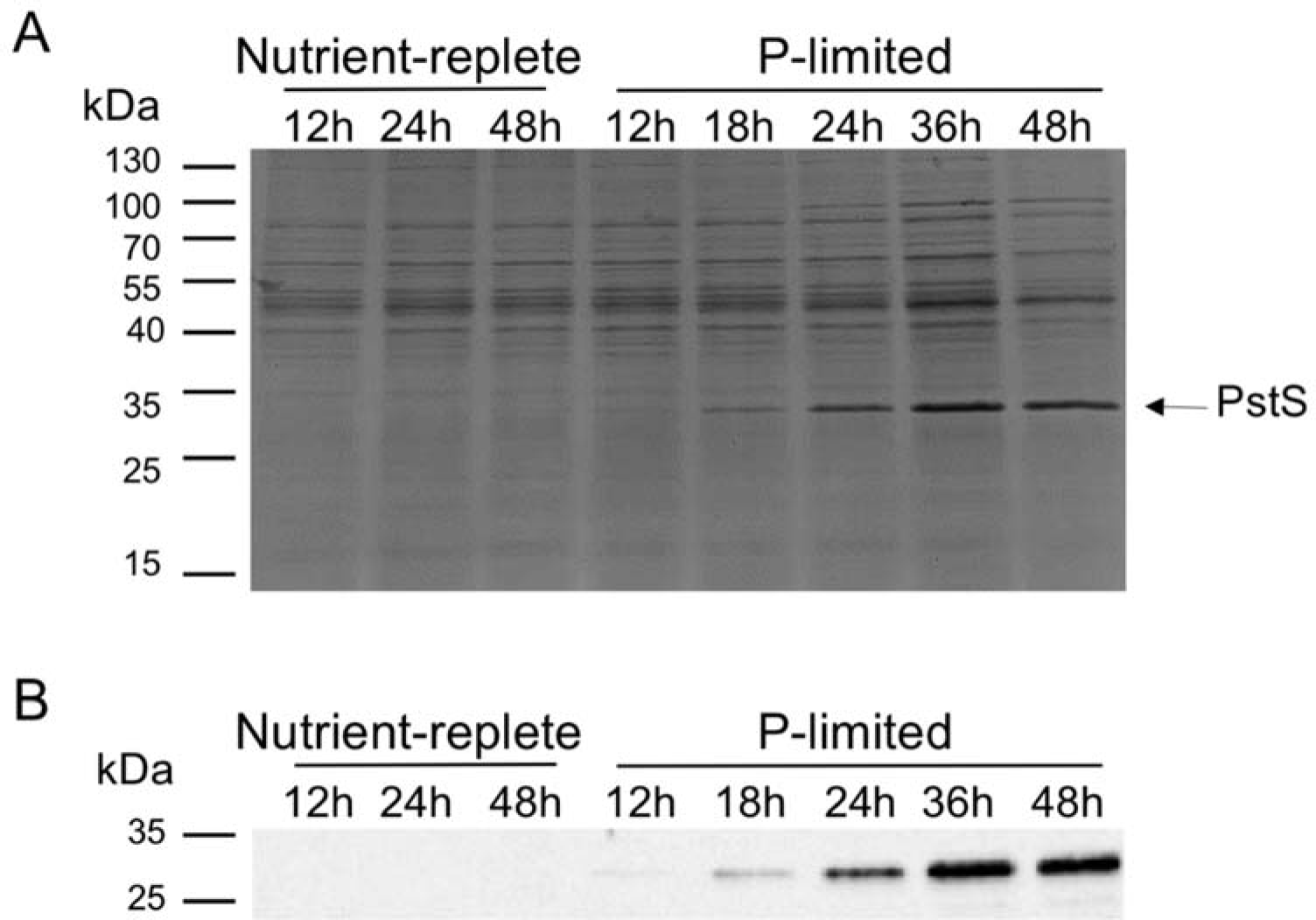
PstS protein abundances of *Prochlorococcus* NATL2A under nutrient-replete and P-limited conditions. **A**. SDS-PAGE. *Prochlorococcus* NATL2A cells were spun down and resuspended in nutrient-replete or P-limited media. Cultures were collected at different time points after resuspension. Total protein from 10^8^ cells was loaded in each lane and separated in SDS-PAGE. The time after resuspension is shown above each lane. Protein sizes are shown on the left of the gel. The arrow indicates the PstS protein band (∼34 kDa). **B**. Western blot. Proteins from replicate SDS-PAGE were transferred to a membrane and probed with the antibody against NATL2A PstS.

**Supplementary Figure 3.**
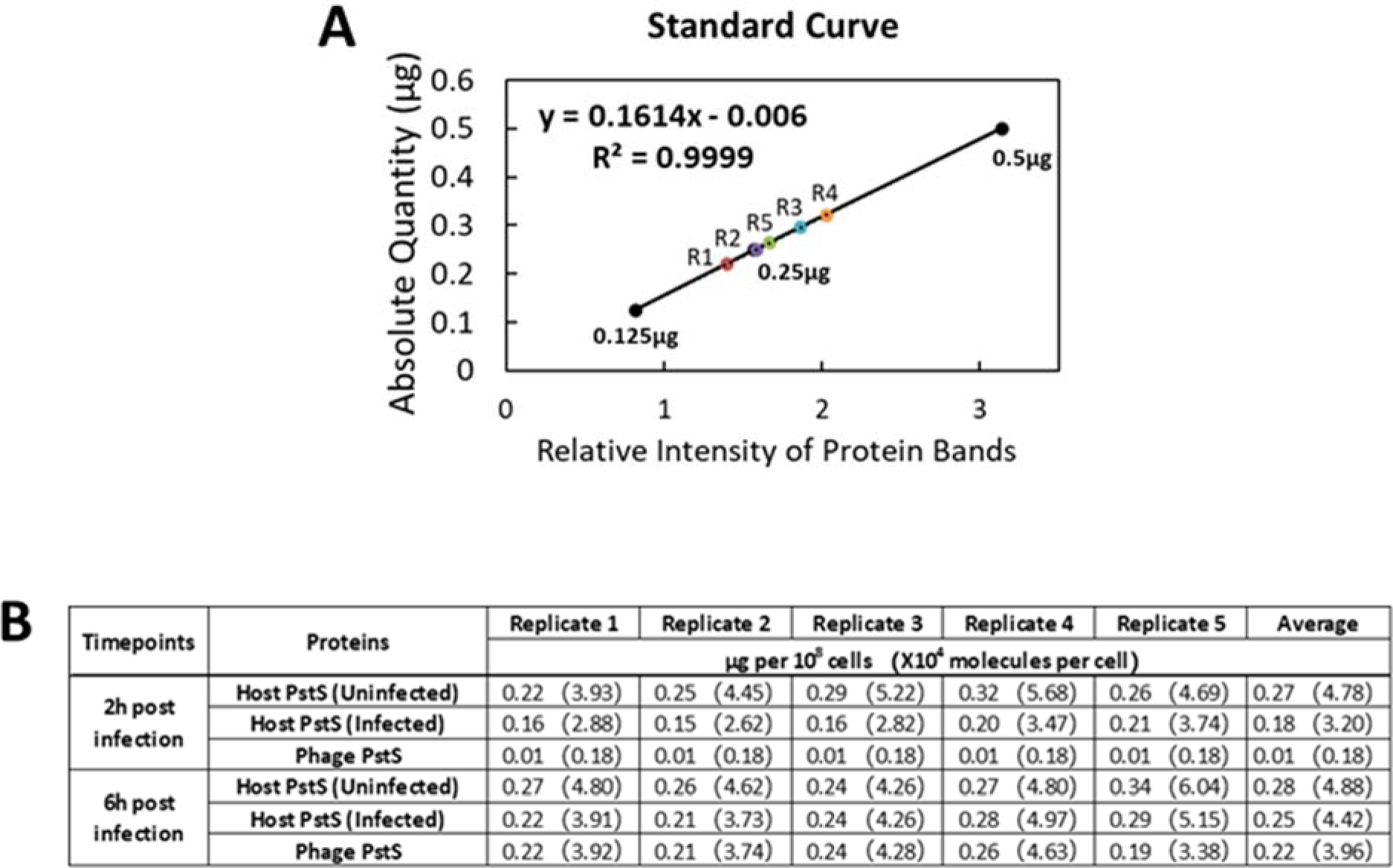
Quantification of PstS proteins in *Prochlorococcus* NATL2A cells using quantitative western blotting. **A**. A representative standard curve generated using the signal volumes of protein bands in the quantitative western blots shown in Figure 2C (top panel). **B**. Based on the standard curve, amounts of PstS proteins per cell were calculated.

**Supplementary Figure 4.**
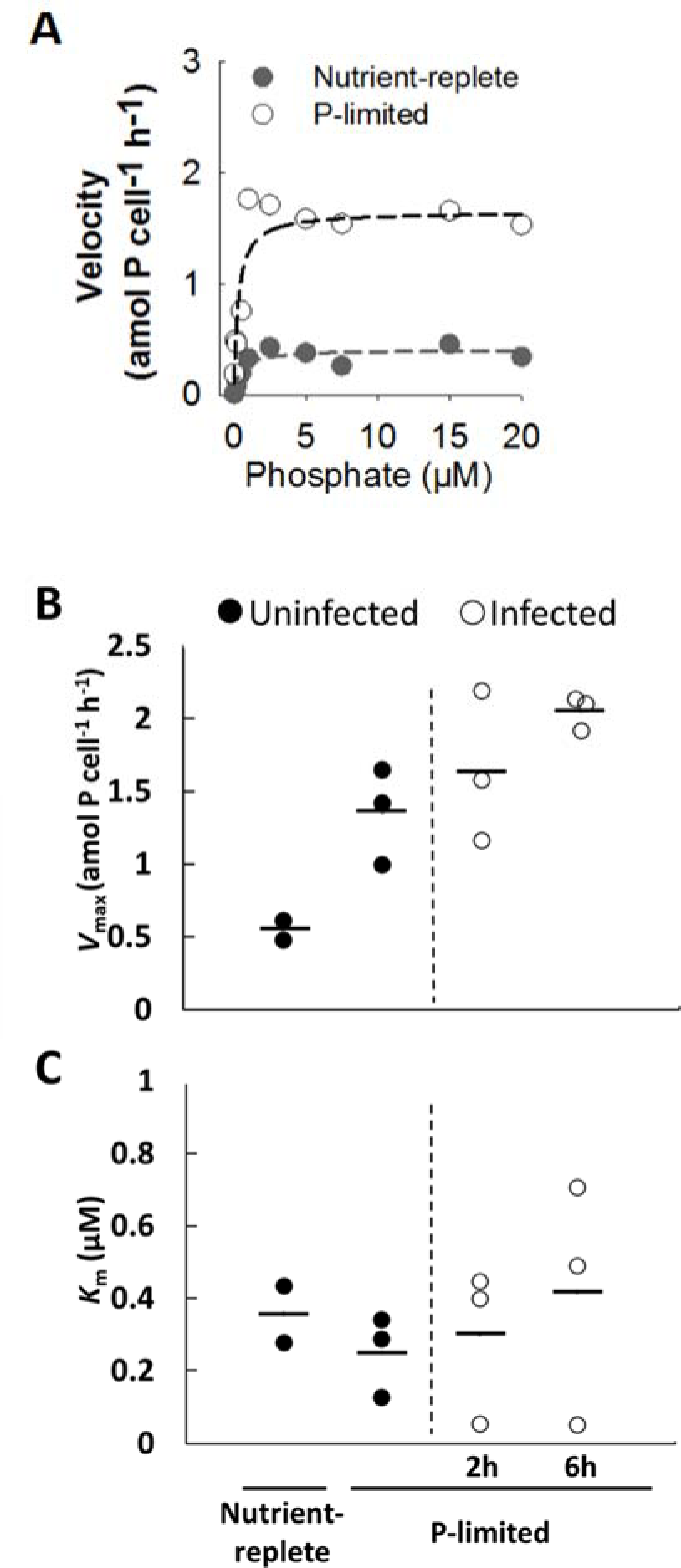
Phosphate uptake of uninfected and infected *Prochlorococcus* NATL2A cells. *Prochlorococcus* NATL2A cells were spun down and resuspended in nutrient-replete or P-limited growth media. At 24 h after resuspension, phosphate uptake velocity of uninfected *Prochlorococcus* NATL2A cells was measured as a function of cold phosphate. At 24 h after resuspension in P-limited growth media, *Prochlorococcus* NATL2A cells were infected by cyanophage P-SSM2 at a phage/host ratio of 3 and phosphate uptake velocity at 2 h and 6 h after infection was measured. **A** represents phosphate uptake curves of uninfected cells. Dashed lines in **A** represent the best fit of a hyperbolic curve. Using the phosphate uptake curves, the maximum uptake velocity (*V*_max_) (**B**) and the Michaelis-Menten constant (*K*_M_) (**C**) of *Prochlorococcus* NATL2A were calculated.

**Supplementary Figure 5.**
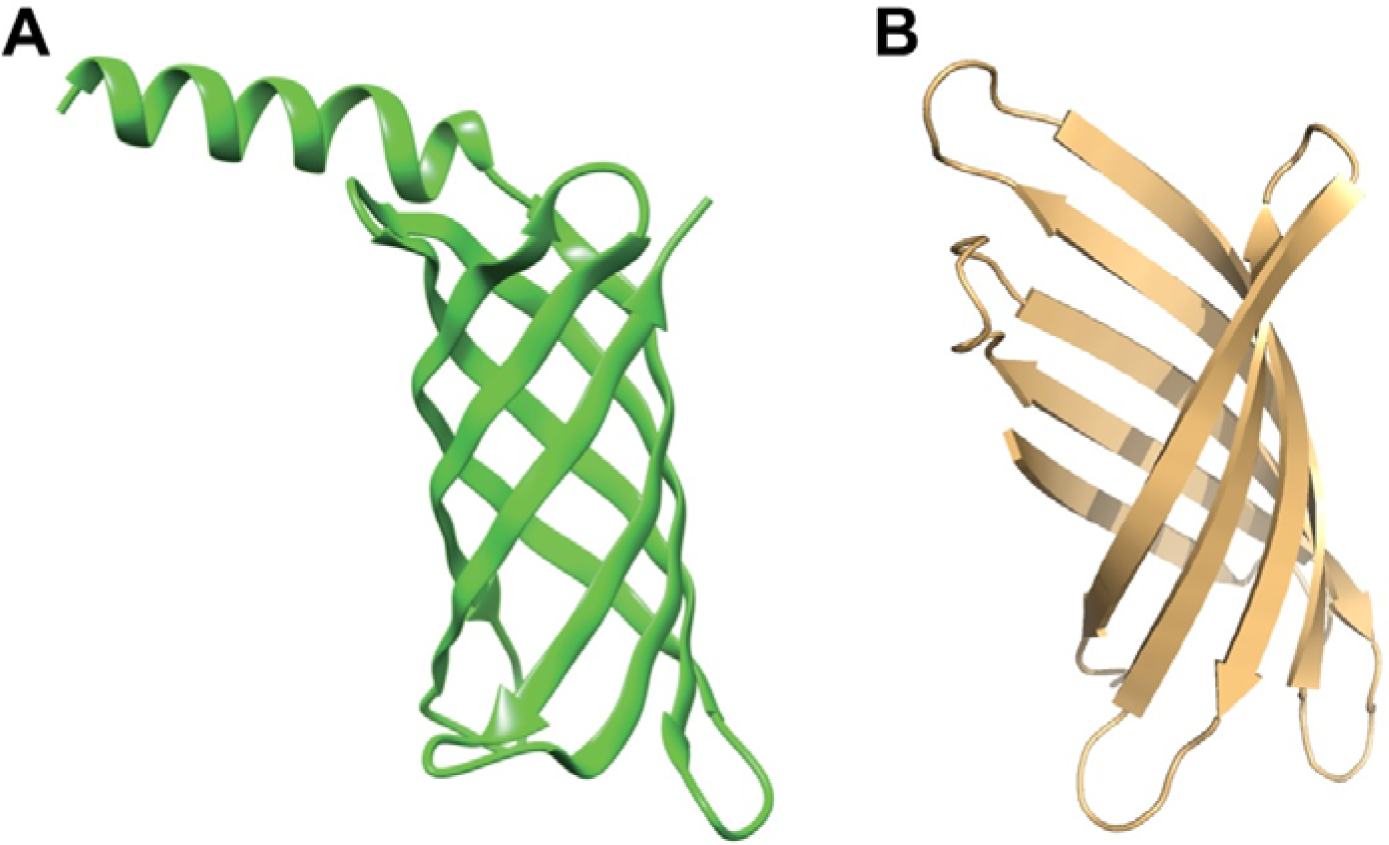
Predicted structures of cyanophage P-SSM2 gp247 Structure of cyanophage gp247 was predicted using two methods. **A**. De novo folding structure was predicted by the tFold server (https://drug.ai.tencent.com/console/en/tfold). The structure with the highest ranking is shown. **B**. Structure of gp247 was predicted by the template-based modeling method via Phyre2 (http://www.sbg.bio.ic.ac.uk/phyre2) based on the three-dimensional structure of the outer membrane porin OprF of *Pseudomonas aeruginosa* (Phyre2 fold library ID: c4rlca; confidence score: 91.3).

**Supplementary Figure 6.**
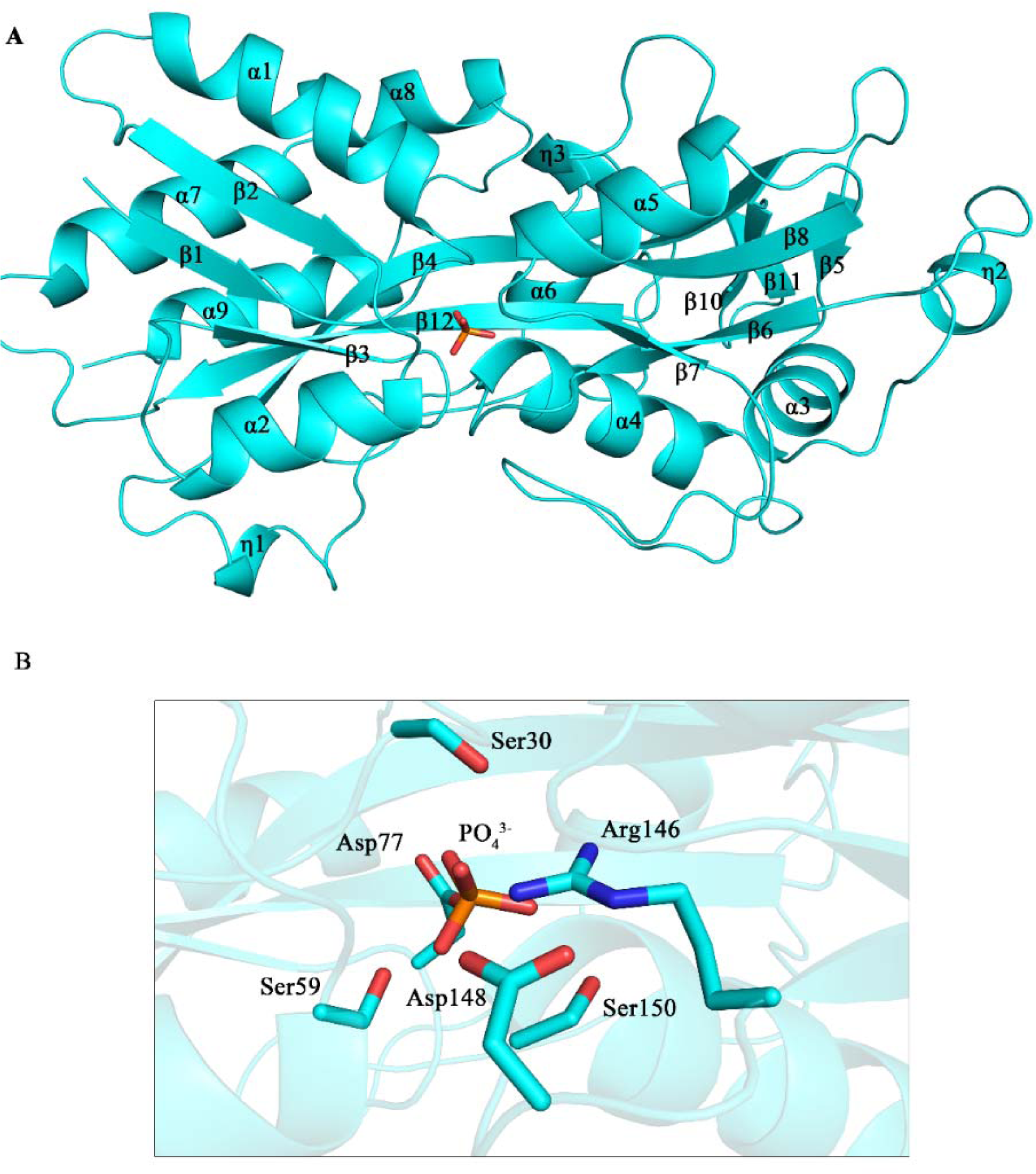
Overall structure of P-SSM2 PstS in complex with PO_4_. **A**. Cartoon representation of the overall structure of the P-SSM2 PstS protein in complex with a PO_4_ molecule (shown in red). Secondary structural elements of PstS are labeled sequentially. **B**. Detailed view of the PO_4_-binding site, with PO_4_-interacting residues shown as sticks.

**Supplementary Figure 7.**
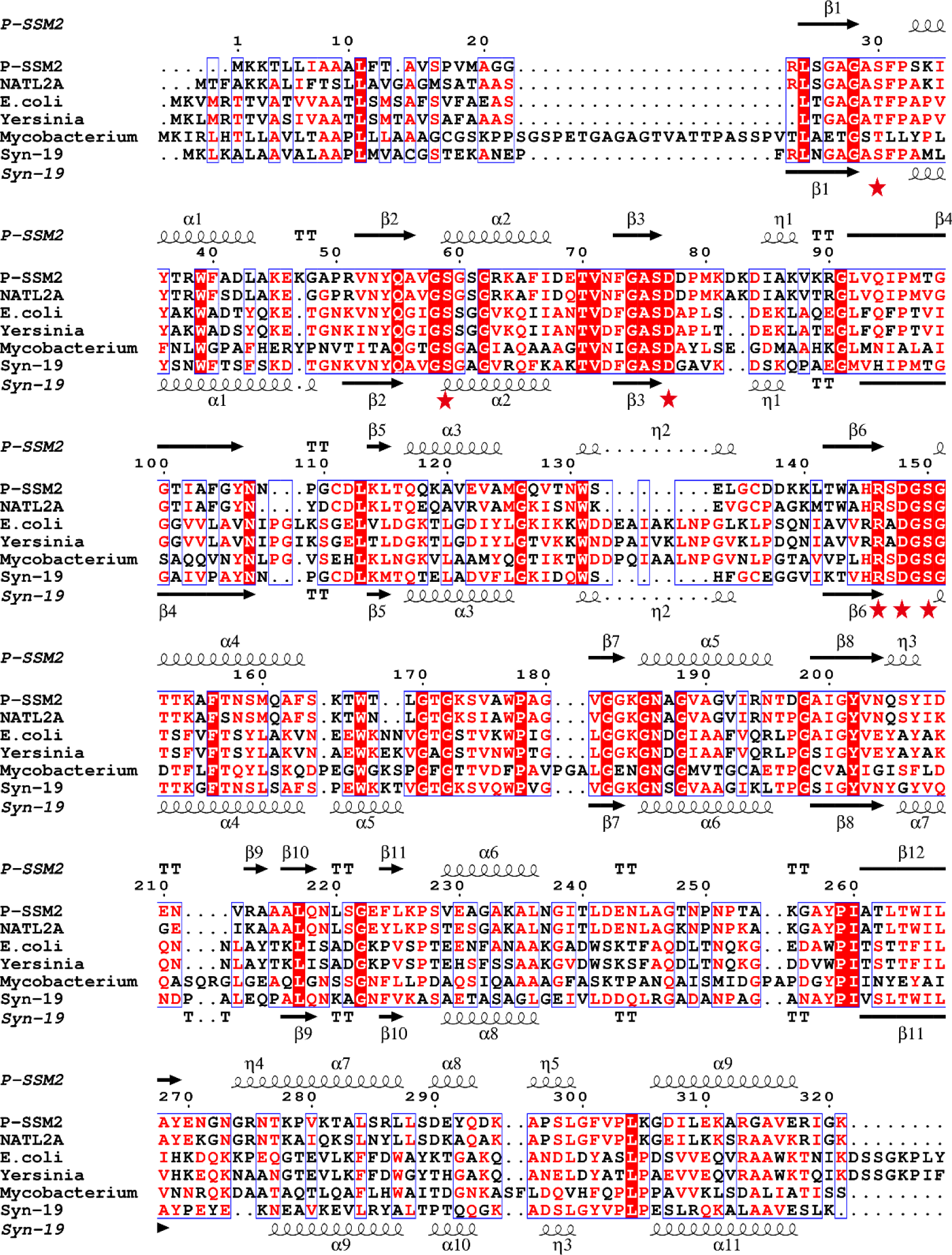
Sequence alignment of PstS proteins. The secondary structural elements of P-SSM2 PstS and Syn19 PstS are shown at the top and bottom of the PstS sequence alignment, respectively. The numbers above the sequence alignment indicate the residue numbers of P-SSM2 PstS. The phosphate-interacting residues of P-SSM2 PstS are marked by red stars. Among these residues, Ser59, Asp77, Arg146, Asp148, and Ser150 are conserved in all the PstS proteins listed here. Ser30 is conserved in cyanophage and cyanobacterial proteins, and is replaced by a chemically similar amino acid threonine in other PstS proteins.

**Supplementary Figure 8.**
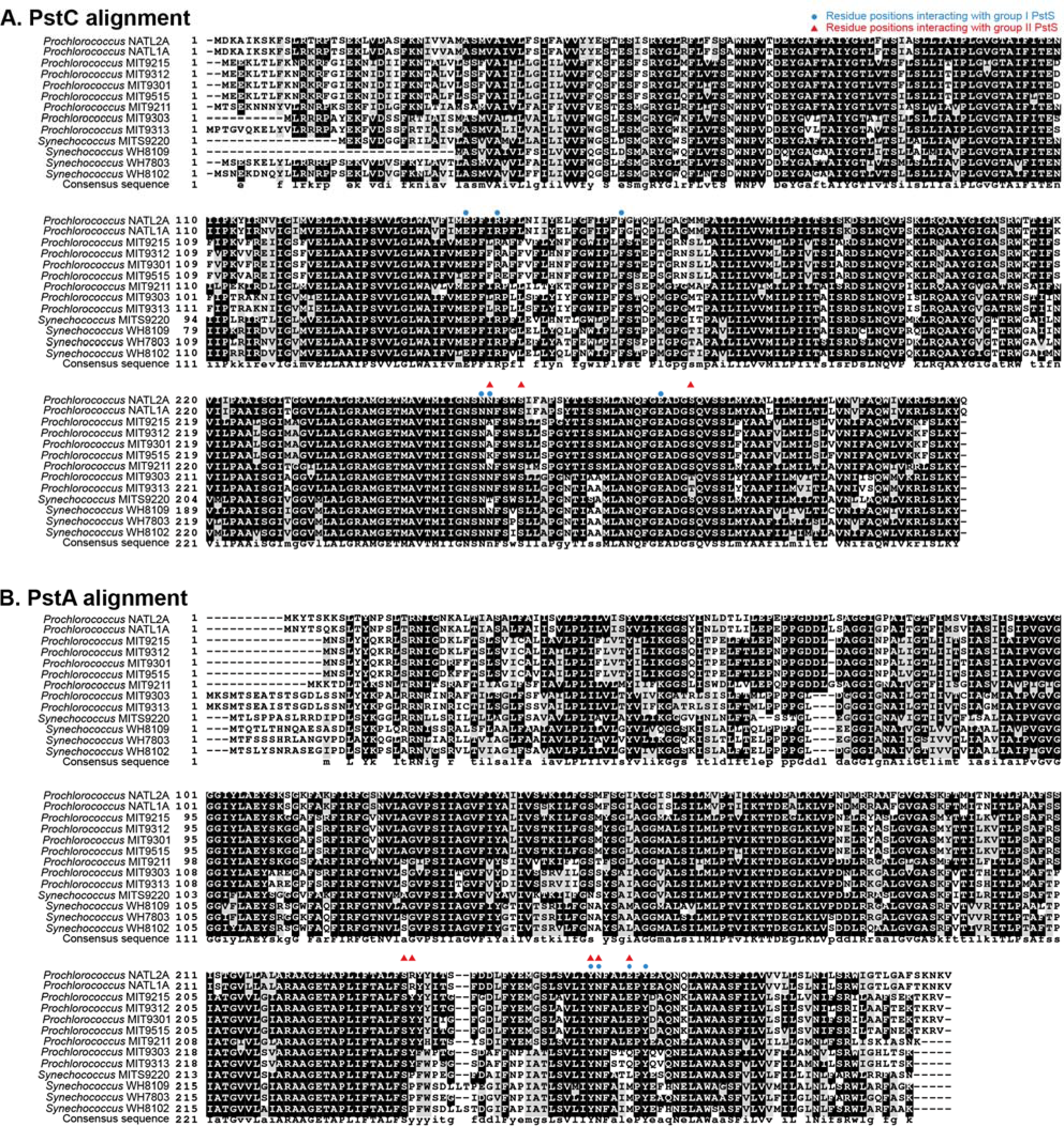
Sequence alignment of cyanobacterial PstC and PstA proteins Protein sequence alignments are shown for cyanobacterial PstC. (**A**) and PstA (**B**). The background shading indicates the degree of conservation, with black for strongly conserved and grey for moderately conserved residues. Putative residues interacting with group I PstS are marked with blue circles and those interacting with group II PstS are marked with red triangles.

**Supplementary Figure 9.**
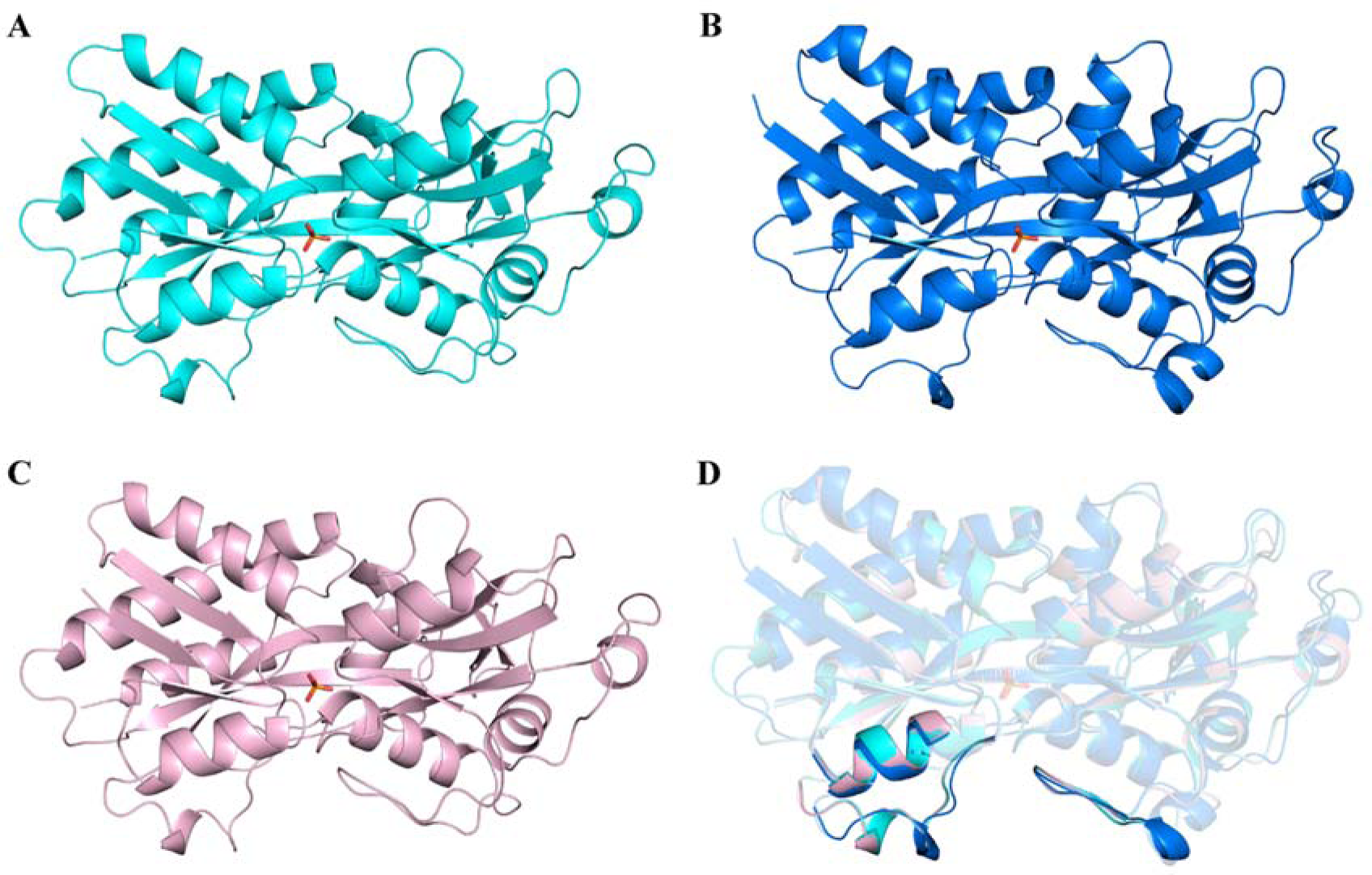
Overall structures of PstS proteins. **A.** The PstS structure of cyanophage P-SSM2 in complex with PO_4_ (red symbol). **B**. The PstS structure of cyanophage Syn19 in complex with PO_4_. **C**. A model of PstS structure of *Prochlorococcus* NATL2A in complex with PO_4_. **D**. Superposition of PstS proteins of P-SSM2 (cyan), Syn19 (blue) and NATL2A (light pink). The segments involved in interaction with PstCA are highlighted.

**Supplementary Figure 10.**
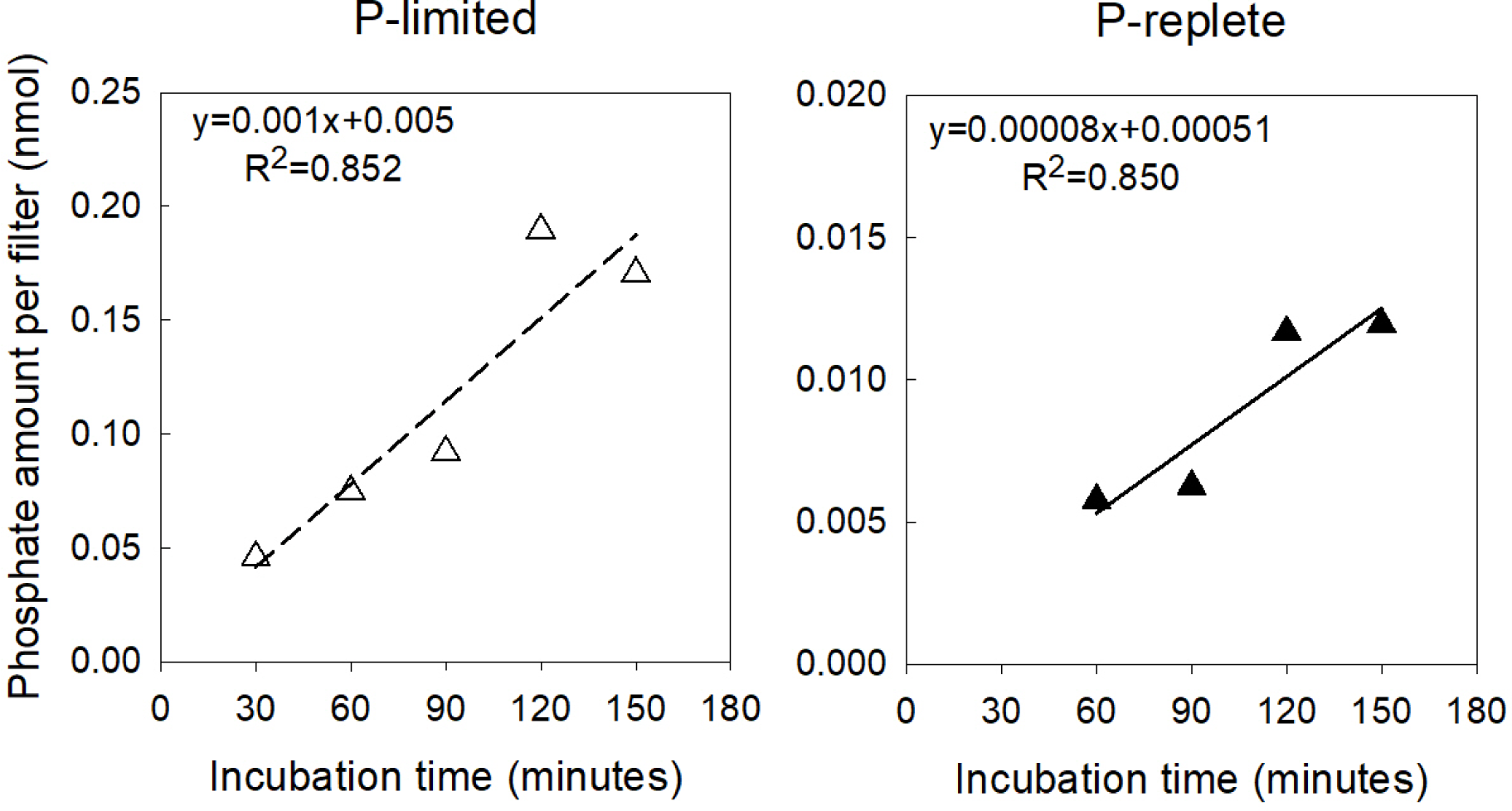
Phosphate uptake by *Prochlorococcus* NATL2A over time. Axenic *Prochlorococcus* NATL2A cells were grown under P-limited (left panel) or P-replete (right panel) conditions. To measure phosphate uptake rates, *Prochlorococcus* cells were pelleted by centrifugation and resuspended with the Pro99 medium without addition of phosphate. Resuspended cells were amended with 10 µM cold NaH_2_PO_4_ and trace amount of ^32^P-labeled orthophosphoric acid (∼1 µCi, Perkin Elmer). Every 30 min, 1 ml culture was filtered through a 0.22 µm polycarbonate filter that was supported by a Whatman GF/F filter. During filtration, the vacuum pressure was ∼100 mm Hg. The phosphate uptake amount by 1 ml culture was estimated by measuring the radioactivity on each filter.

